# Enhancing Immune Cell Activation through Cold Atmospheric Plasma: Disruption of PD-1/PD-L1/PD-L2 Immune Checkpoints via Molecular Dynamics

**DOI:** 10.1101/2025.01.23.634633

**Authors:** Rasulbek Mashalov, Zulfizar Toshpulatova, Zhitong Chen, Jamoliddin Razzokov

## Abstract

The PD-1/PD-L1/PD-L2 immune checkpoint plays a critical role in regulating immune responses, and its dysfunction is implicated in immune evasion by cancer cells. Cold atmospheric plasma (CAP) has emerged as a promising cancer therapeutic modality with the potential to modulate immune checkpoints. This study employs molecular dynamics (MD) simulations to investigate the impact of CAP-induced oxidation on the interactions between PD-1 and its ligands, PD-L1 and PD-L2. We simulated the PD-1/PD-L1 and PD-1/PD-L2 complexes under different oxidation levels. Key residues within the interaction site of the ligands are modified using Vienna PTM 2.0 online server. Umbrella sampling and other MD analyses revealed that increasing oxidation levels leads to weaken the binding affinity between PD-1 and both PD-L1 and PD-L2. These findings suggest that CAP may offer a novel strategy for enhancing anti-tumor immunity. This computational study provides valuable insights into the molecular mechanisms underlying CAP’s effects on immune regulation and highlights its potential for cancer immunotherapy.

## Introduction

Cancer remains a major global cause of mortality, with millions of new diagnoses made each year [1]. Treatments such as surgery, chemotherapy, and radiation have certainly saved many lives, however they often come with some drawbacks. These treatments may cause side effects, for example damaging healthy tissues, leading to toxicity throughout the body, and negatively affecting a patient’s overall life quality [2-5]. These challenges show the need for new, more accurate, and less invasive treatment methods. Among the emerging alternative approaches, the use of ionized gas in the form of cold atmospheric plasma (CAP), also called non-thermal plasma, is gaining considerable attention for its electively target cancer cells while preserving normal tissues [6, 7]. It can be generated at room temperature and atmospheric pressure using different plasma sources, including dielectric barrier discharges (DBDs), plasma jets, and micro plasma devices, which allow for the delivery of bioactive species while minimizing thermal damage to surrounding tissues [8-10].

Some studies have demonstrated the potential of CAP in inducing apoptosis and halting cancer cell proliferation across different cancer types. For example, Wang et al. showed how CAP induces apoptosis and inhibits proliferation in colon and lung cancer cells [11]. While Weiss et al. suggested that CAP treatment reduces cell growth and induces apoptosis in prostate cancer cells [12]. One of the important characteristics of CAP is its selective action, targeting cancerous cells while sparing normal ones [13]. With this feature the tumor cell can be targeted and exposed to flow of reactive oxygen and nitrogen species (RONS), generated by CAP, which eventually make the tumor cell more vulnerable to apoptosis induced by immune cell recognition [8, 9, 14, 15].

Additionally, CAP has been studied for its potential to target immune checkpoints proteins (ICPs) on cancer cells [16-19]. These proteins are considered as transmembrane receptors with variety downstream signaling pathways that branch out upon ligand-receptor interaction and are expressed on multiple types of immune cells, and their expression can vary within a single cell type [20-30]. This natural variation in expression patterns complicates the targeting of specific effector immune cells in cancer treatment [31-33]. Among the immune checkpoint proteins, the programmed cell death protein 1 (PD-1) and its ligands, PD-L1 and PD-L2, are key players in the immune system’s regulation of cancer immunity [34]. PD-1 is primarily expressed on T cells, and its interaction with PD-L1 or PD-L2, often upregulated on tumor cells, suppresses T cell activity, allowing tumor cells to evade immune detection and destruction [35, 36]. CAP-generated RONS can oxidize proteins such as PD-L1, potentially disrupting its binding with PD-1. This disruption could enhance immune recognition of tumor cells and promote their elimination by T cells [37]. This research investigates the effect of CAP on ICPs. To achieve this, we simulated the oxidation of vulnerable residues, mimicking the impact of CAP treatment on ICPs. MD simulations were conducted, along with the umbrella sampling (US) method, to evaluate the binding affinity between PD-1 and its ligands, PD-L1 and PD-L2. Our findings highlight that CAP-induced oxidation may weaken these interactions, leading to immune recognition and facilitating tumor cell elimination.

## Methods

In our study we used GPU-accelerated GROMACS 5.1 package to conduct MD simulations on PD-1 and its ligands PD-L1/PD-L2 [38-40]. Initial structures of the PD-1 ligands, PD-L1 and PD-L2, were obtained from the Protein Data Bank (PDB), with PDB IDs 4ZQK and 6UMT, respectively [41]. For each ligand complex, three separate models were created: native PD-L1/PD-L2-PD-1, moderately oxidized PD-*L*1_*OX*1_ /PD-*L*2_*OX*1_–PD-1, and highly oxidized PD-*L*1_*OX*2_/PD-*L*2_*OX*2_-PD-1. We utilized Vienna-PTM 2.0 web server to modify specific residues in ligand complexes [42]. Furthermore, residues in PD-L1 and PD-L2 were chosen based on computational analyses of the solvent-accessible surface area (SASA) of various amino acids (AAs) and experimental findings [43]. Selected residues were identified as having higher vulnerability of interacting with plasma-generated reactive oxygen and nitrogen species (RONS). The GROMOS 54a7 force field was employed to generate the topologies. This force field was chosen because it has been widely used for simulating protein-ligand interactions. Studies have shown that GROMOS 54a7 accurately captures the conformational dynamics of immune checkpoint interactions, making it suitable for modeling PD-1/PD-L1 interactions [44, 45]. To make simulation system, a dodecahedron-shaped box is created, as it is considered as a cost-efficient and may shorten the simulation time [46].

The box is then solvated with water SPC water model for both the PD-L1 and PD-L2 complexes, along with 0.1 M Na+ and Cl−, which are subsequently used to electrically neutralize the system [47, 48]. Visualizations of obtained results are generated using PYMOL and VMD tools [49, 50]. Before equilibration, the systems were energy minimized using the steepest descent algorithm. Subsequently, the systems were equilibrated for 10 ps in the NVT ensemble (i.e., a system with constant number of particles N, volume V, and temperature T), with position restraints on the heavy atoms of the proteins. Then, another 10 ps equilibration was performed in the NPT ensemble (i.e., a system with constant number of particles N, pressure p, and temperature T) using weak coupling thermo– and barostats. Finally, 500 ns production runs were carried out, again using the NPT ensemble, but with the position restraints removed, and employing the Nose-Hoover thermostat and the isotropic Parrinello-Rahman barostat. All simulations were performed at 310 K and 1 bar, with a 1.4 nm cut-off for Coulomb and van der Waals interactions. The trajectory from the production runs was used to collect data, i.e., for the calculation of the root mean square deviation (RMSD) and the root mean square fluctuations of PD-퐿1 /PD-퐿2 –PD-1 and PD-퐿1 /PD-퐿2 –PD-1 complexes.

**Table 1.** Amino acids (AA) involved in the creation of the oxidized PD-L1 protein.

| *AA in native PDL-1* | *Modified AA in PDL-1<sub>ox1-ox2</sub>* |
| --- | --- |
| **Methionine (Met)- 36,59,115** | Methionine sulfoxide |
| **Tyrosine (Tyr)- 28,32,56,81,112,118,123** | 3,4- dihydroxyphenylalanine |
| **Phenylalanine (Phe)-19,42,67** | 3,4- dihydroxyphenylalanine |
| **Lysine (Lys)-124** | Allysine |
| **Cysteine (Cys)-40,114** | Cysteic acid |
| **Histidine (His)-69,78** | 2- oxo-histidine |
| **Tryptophan (Trp)-57** | 6- hydroxytryptophan |

To create the oxidized structures of PD-L1, referred to as PD-L1ox1 and PD-L1ox2, 8% (ox1) and 17% (ox2) of the residues were oxidized. The oxidized forms of PD-L1 and PD-L2 were generated using the Vienna-PTM 2.0 [42] web server by substituting native residues in PD-L1/PD-L2 with their oxidized counterparts.

**Table 2.**
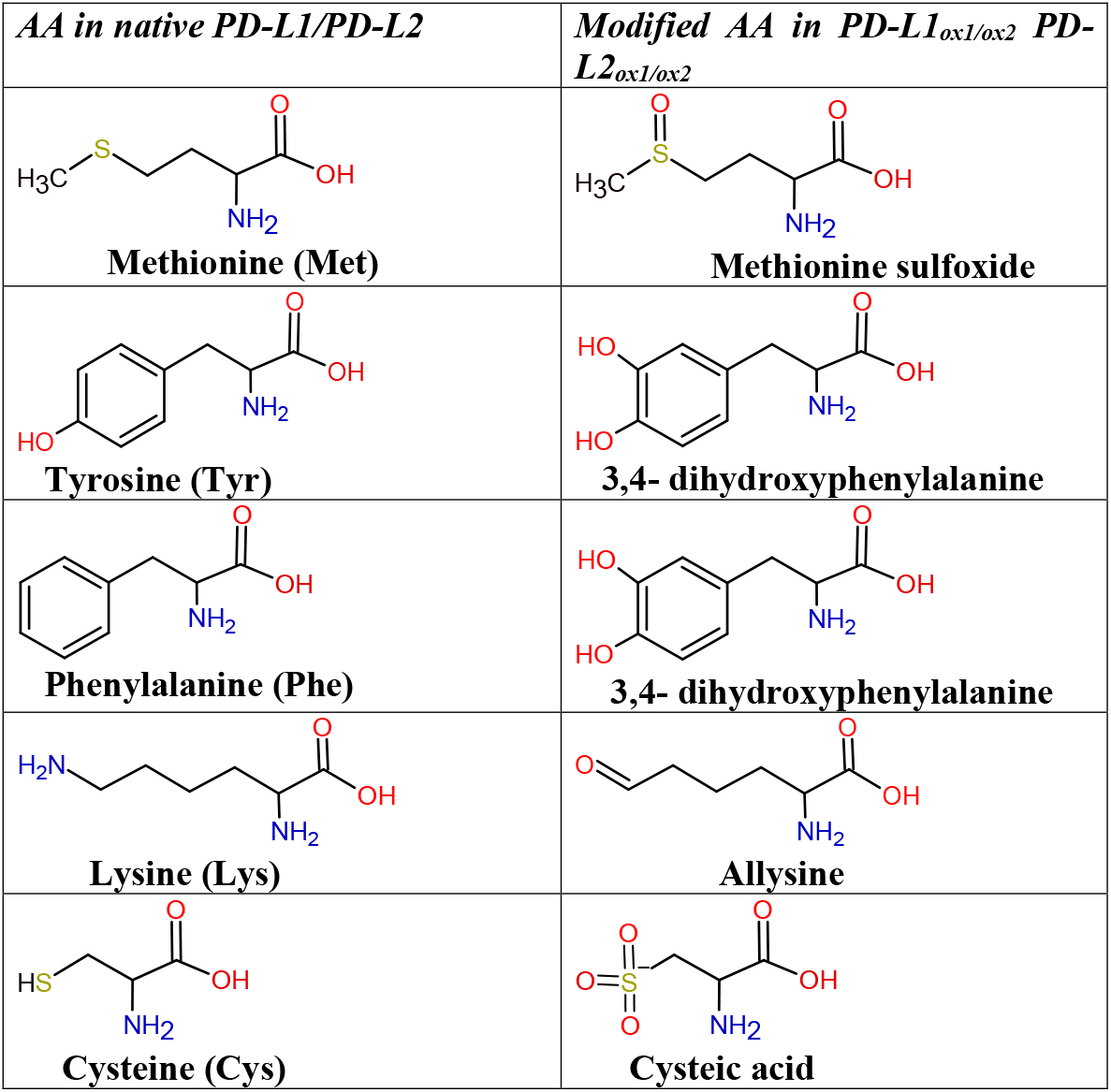
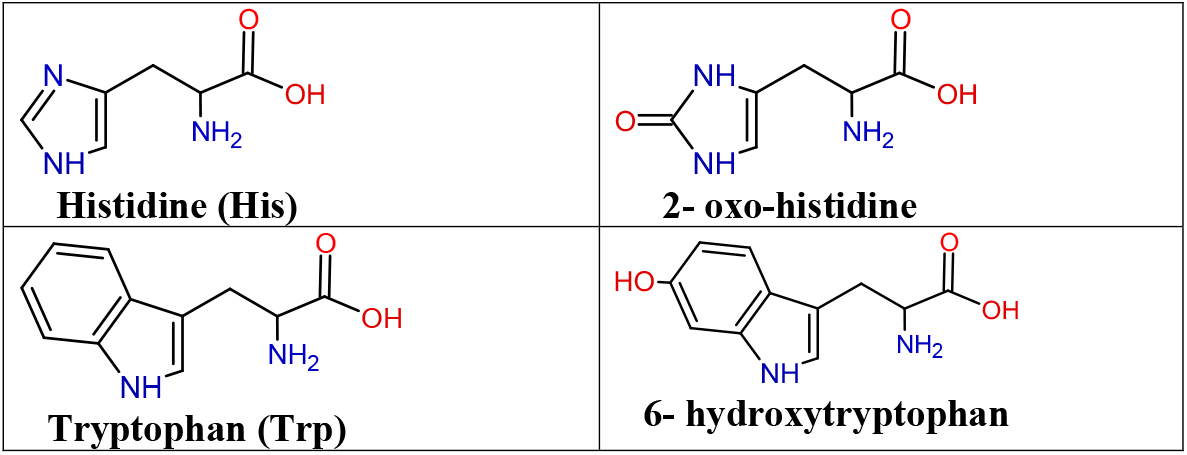
Chemical structures of original and modified AAs in PD-L1/PD-L2 proteins.

Before equilibration, the systems were energy minimized using the steepest descent algorithm. Subsequently, the systems were equilibrated for 10 ps in the NVT ensemble (i.e., a system with constant number of particles N, volume V, and temperature T), with position restraints on the heavy atoms of the proteins. Then, another 10 ps equilibration was performed in the NPT ensemble (i.e., a system with constant number of particles N, pressure p, and temperature T) using weak coupling thermo– and barostats. Finally, 500 ns production runs were carried out, again using the NPT ensemble, but with the position restraints removed, and employing the Nose-Hoover thermostat and the isotropic Parrinello-Rahman barostat. All simulations were performed at 310 K and 1 bar, with a 1.4 nm cut-off for Coulomb and van der Waals interactions. The trajectory from the production runs was used to collect data, i.e., for the calculation of the root mean square deviation (RMSD) and the distance between salt bridges of the PD-L1/PD-L2/PD-L1ox1/PD-L2ox1/PD-L1ox2/PD-L2ox2 and PD-1 complexes. The PyMOL molecular graphics tool was used to prepare the images presented in this paper.

From the final 50 ns of each 500 ns production trajectory, systems were extracted at 450 ns for both native and oxidized complexes, resulting in six model systems in total. These models were then utilized in umbrella sampling (US) simulations. For the US simulations, 35 windows were used, spaced 0.1 nm apart, which were derived from pulling simulations of PD-L1/PD-L1ox1/ox2 and PD-L2/PD-L2ox1/ox2. During the pulling simulations, native and oxidized forms of PD-L1 and PD-L2 were restrained and served as references, while PD-1 was pulled along the z-axis. Additionally, PD-1’s movement in the xy-plane was restrained using flat-bottom position restraints with a radius of 0.1 nm and a force constant of 500 kJ/(mol·nm^2^).

The pulling simulations applied a spring constant of 1000 kJ/(mol·nm^2^) and a pulling rate of 0.001 nm/ps, which resulted in the dissociation of PD-1 from native and oxidized forms of PD-L1 and PD-L2. Each umbrella window was then subjected to 20 ns of US simulations, with the last 17 ns used for data collection after a 3 ns relaxation period. The weighted histogram analysis method (WHAM) was employed to calculate the potential of mean force (PMF) or free energy profiles (FEPs) for PD-1 dissociation from native and oxidized PD-L1 and PD-L2 complexes. Errors in the FEPs were estimated using the bootstrapping method. Overall, 210 US simulations (35 windows × 1 trajectory × 6 systems) were conducted to derive the final FEPs.

## Results and Discussion

Protein oxidation is a well-documented phenomenon that increases structural flexibility, often leading to significant conformational changes. In this study, we investigated the effects of oxidation on the structural and functional properties of PD-L1 and PD-L2. RMSD (root mean square deviation) calculations of the backbone atoms for both native and oxidized forms revealed that oxidation induces higher fluctuations in the protein structures (Figure 1). These increased fluctuations are indicative of enhanced molecular flexibility, which in turn facilitates conformational changes. The most notable changes were observed in the oxidized forms, PD-L1ox1/ox2 and PD-L2ox1/ox2, highlighting the structural destabilization caused by oxidative modifications.

**Figure 1:**
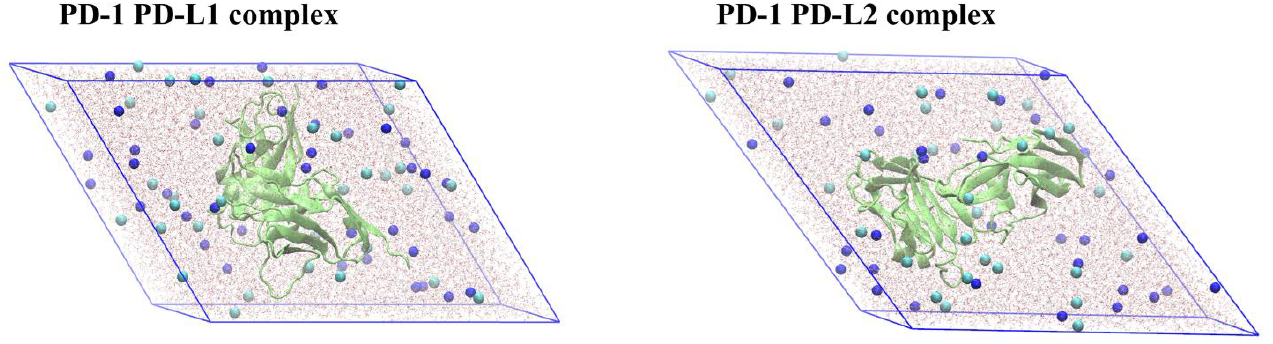
Initial states of the complexes before the start of the simulation.

**Figure 2.**
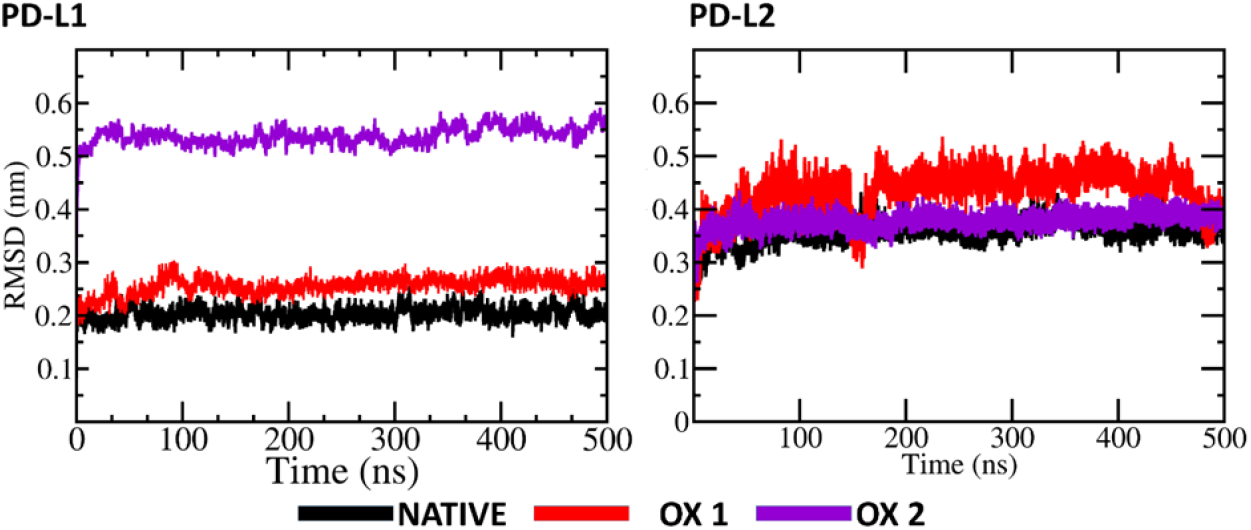
The root mean square deviation (RMSD) of the backbone atoms for both native and oxidized structures PD-L1/PD-L2 and oxidized structures.

To further understand the functional implications of these structural changes, we carried out molecular dynamics (MD) simulations to analyze the interactions of native and oxidized PD-L1 and PD-L2 with their respective binding partners. These simulations revealed that NPT-induced oxidation weakens the binding affinity of PD-L1 and PD-L2 to PD-1 protein. This reduction in binding affinity is attributed to structural changes at the binding interface, likely caused by the disruption of stabilizing interactions such as hydrogen bonds and van der Waals forces.

The alignment of PD-L1–PD-1 and PD-L1ox–PD-1 systems provided further insights into the structural effects of oxidation. A clear shift in the loops located at the binding sites of PD-L1ox and PD-1 was observed (Figure 2). These shifts are a direct consequence of the conformational changes in PD-L1ox, driven by oxidation-induced alterations in the protein’s secondary and tertiary structures. These loop shifts may disrupt the precise spatial arrangement required for optimal binding, leading to a decrease in binding affinity. The impact of these structural changes is consistent with the calculated potential of mean force (PMF) profiles, which indicate a less favorable interaction between oxidized PD-L1/PD-L2 and PD-1 compared to their native counterparts.

**Figure 2.**
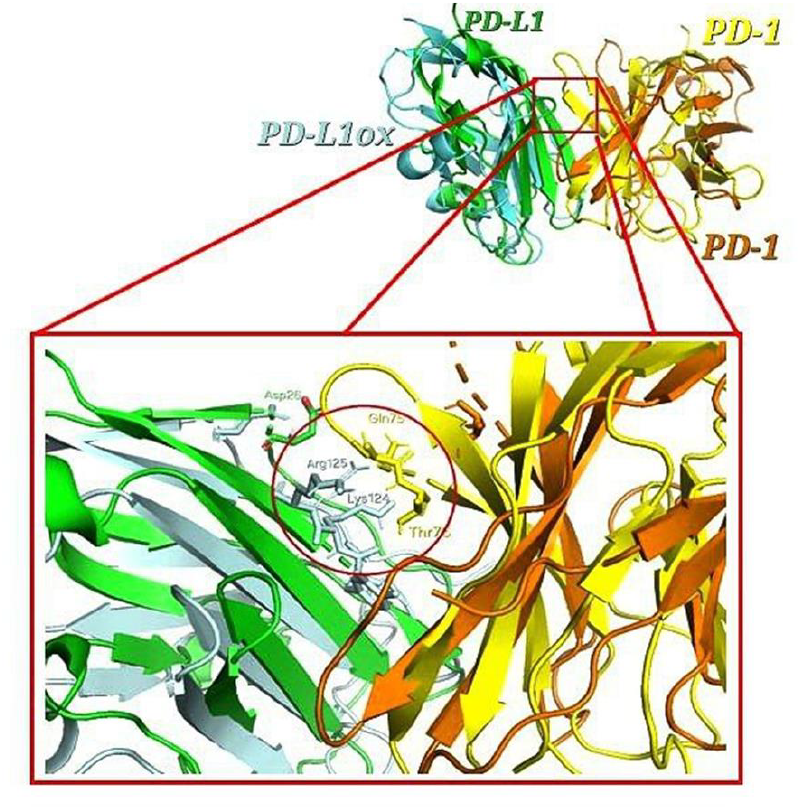
The interaction of PD-1 and PD-L1 proteins.

The implications of these findings extend beyond structural considerations, as they also affect the immunological roles of these proteins. Oxidation-induced modifications may impair the ability of PD-L1 and PD-L2 to interact effectively with PD-1, a key immune checkpoint receptor. This could potentially alter the immune evasion mechanisms employed by cells expressing these proteins, such as tumor cells. Furthermore, reduced binding to PD-1 suggests that oxidation may also affect the therapeutic targeting of PD-L1/PD-L2 in immunotherapy applications.

**Figure 3.**
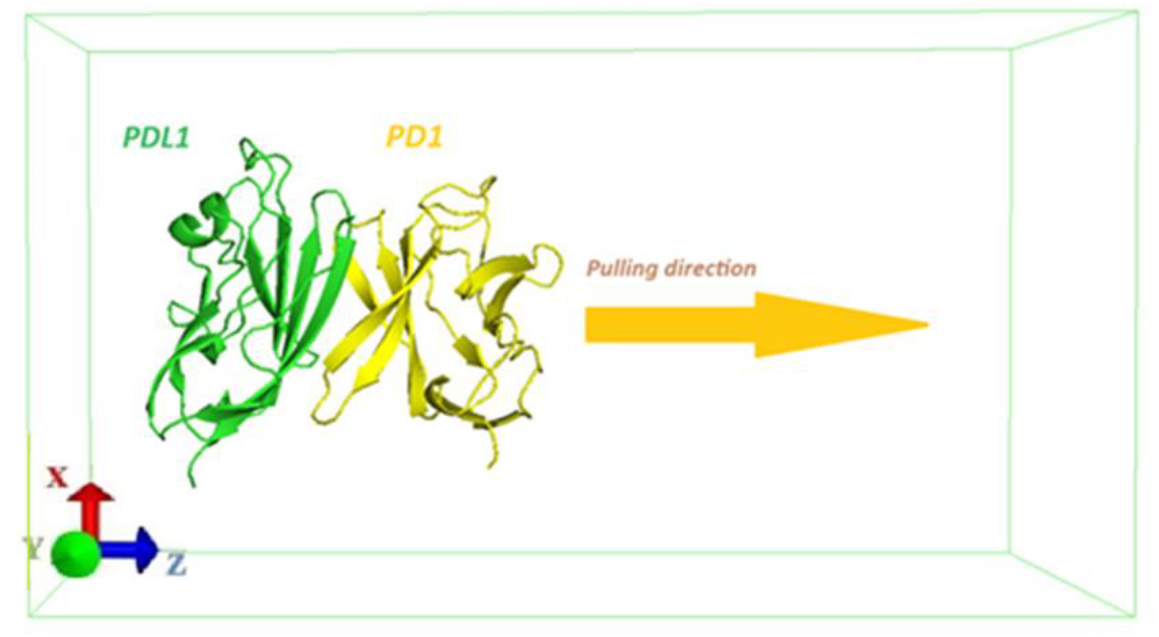
PD-1 was pulled against native and oxidized PD-L1 structures.

**Figure 4.**
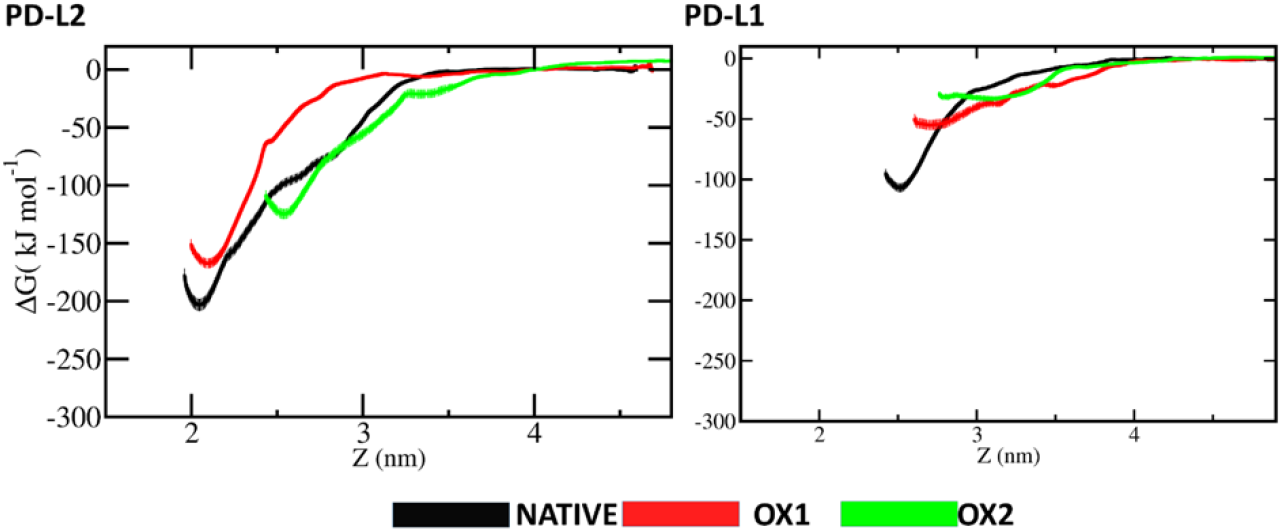
Free energy profiles (FEPs) of the native (black), oxidized (red) and highly oxidized (green) PD-L1/PD-L2 structures dissociated from PD-1, indicate the dissociation energy.

## Conclusions

In summary, our results demonstrate that protein oxidation leads to significant structural changes in PD-L1 and PD-L2, particularly in regions critical for binding. These conformational alterations result in reduced binding affinities to PD-1 and therapeutic antibodies, highlighting the critical role of oxidative modifications in regulating protein function. This study provides valuable insights into the molecular mechanisms by which oxidative stress can modulate immune checkpoint interactions. Such modifications may serve as a natural mechanism to enhance immune activation or, conversely, as a contributor to immune evasion in pathological conditions such as cancer and chronic inflammation. Understanding these dynamics could offer novel perspectives on how oxidative stress influences immune regulation.

Furthermore, the findings underscore the potential of targeting oxidative pathways in therapeutic interventions aimed at modulating immune checkpoint activity. For instance, leveraging oxidative processes through treatments like cold atmospheric plasma may provide a promising approach to disrupt immune checkpoint interactions and boost immune responses in cancer therapy. Our molecular dynamics simulations also pave the way for future studies to explore other immune-related proteins impacted by oxidative modifications. Investigating these effects in a broader biological context will be crucial for translating these findings into clinical applications. Overall, this research not only sheds light on the complex interplay between oxidative stress and immune regulation but also highlights its potential for driving innovation in immunotherapy strategies.

## Acknowledgements

Z.C. acknowledges funding from the National Key Research and Development Program of China (Grant No. 2022YFE0126000). J.R. gratefully acknowledges financial support from the Agency for Innovative Development of the Republic of Uzbekistan (Grant No. AL-59-21122141).

## Notes

### Competing Interest Statement

The authors have declared no competing interest.

## References

1. Siegel, R.L., et al., Cancer statistics, 2021. CA: a cancer journal for clinicians, 2021. 71(1): p. 7–33.

2. Wyld, L., R.A. Audisio, and G.J. Poston, The evolution of cancer surgery and future perspectives. Nature reviews Clinical oncology, 2015. 12(2): p. 115–124.

3. Amjad, M.T., A. Chidharla, and A. Kasi, Cancer chemotherapy. 2020.

4. Baskar, R., et al., Cancer and radiation therapy: current advances and future directions. International journal of medical sciences, 2012. 9(3): p. 193.

5. Richardson, J.L., G. Marks, and A. Levine, The influence of symptoms of disease and side effects of treatment on compliance with cancer therapy. Journal of Clinical Oncology, 1988. 6(11): p. 1746–1752.

6. Dai, X., et al., The emerging role of gas plasma in oncotherapy. Trends in biotechnology, 2018. 36(11): p. 1183–1198.

7. Keidar, M. Cold atmosphere plasma in cancer therapy. in APS Division of Plasma Physics Meeting Abstracts. 2012.

8. Lu, X., et al., Reactive species in non-equilibrium atmospheric-pressure plasmas: Generation, transport, and biological effects. Physics Reports, 2016. 630: p. 1–84.

9. Lin, A., et al., Nanosecond-pulsed DBD plasma-generated reactive oxygen species trigger immunogenic cell death in A549 lung carcinoma cells through intracellular oxidative stress. International journal of molecular sciences, 2017. 18(5): p. 966.

10. Khanikar, R.R., et al., Cold atmospheric pressure plasma technology for biomedical application. Plasma Sci. Technol, 2021.

11. Wang, Y., et al., Cold atmospheric plasma induces apoptosis in human colon and lung cancer cells through modulating mitochondrial pathway. Frontiers in Cell and Developmental Biology, 2022. 10: p. 915785.

12. Weiss, M., et al., Cold atmospheric plasma treatment induces anti-proliferative effects in prostate cancer cells by redox and apoptotic signaling pathways. PloS one, 2015. 10(7): p. e0130350.

13. Semmler, M.L., et al., Molecular mechanisms of the efficacy of cold atmospheric pressure plasma (CAP) in cancer treatment. Cancers, 2020. 12(2): p. 269.

14. Babajani, A., et al., Reactive oxygen species from non-thermal gas plasma (CAP): implication for targeting cancer stem cells. Cancer Cell International, 2024. 24(1): p. 344.

15. Živanić, M., et al., Current state of cold atmospheric plasma and cancer-immunity cycle: therapeutic relevance and overcoming clinical limitations using hydrogels. Advanced Science, 2023. 10(8): p. 2205803.

16. Lin, A., et al., Oxidation of innate immune checkpoint CD47 on cancer cells with non-thermal plasma. Cancers, 2021. 13(3): p. 579.

17. Wei, J., et al., Current trends in sensitizing immune checkpoint inhibitors for cancer treatment. Molecular Cancer, 2024. 23: p. 279.

18. Heirman, P., et al., Effect of plasma-induced oxidation on NK cell immune checkpoint ligands: A computational-experimental approach. Redox Biology, 2024. 77: p. 103381.

19. Wang, Y., et al., Cold atmospheric plasma sensitizes head and neck cancer to chemotherapy and immune checkpoint blockade therapy. Redox Biology, 2024. 69: p. 102991.

20. Freeman, G.J., et al., Engagement of the PD-1 immunoinhibitory receptor by a novel B7 family member leads to negative regulation of lymphocyte activation. The Journal of experimental medicine, 2000. 192(7): p. 1027–1034.

21. Philips, E.A., et al., Transmembrane domain–driven PD-1 dimers mediate T cell inhibition. Science immunology, 2024. 9(93): p. eade6256.

22. Xu-Monette, Z.Y., J. Zhou, and K.H. Young, PD-1 expression and clinical PD-1 blockade in B-cell lymphomas. Blood, The Journal of the American Society of Hematology, 2018. 131(1): p. 68–83.

23. Keir, M.E., et al., PD-1 and its ligands in tolerance and immunity. Annu. Rev. Immunol., 2008. 26(1): p. 677–704.

24. Lu, Y., et al., Cold atmospheric plasma for cancer treatment: molecular and immunological mechanisms. IEEE Transactions on Radiation and Plasma Medical Sciences, 2022. 6(8): p. 916–927.

25. Han, X., et al., Local and targeted delivery of immune checkpoint blockade therapeutics. Accounts of chemical research, 2020. 53(11): p. 2521–2533.

26. Ishida, Y., et al., Induced expression of PD-1, a novel member of the immunoglobulin gene superfamily, upon programmed cell death. The EMBO journal, 1992. 11(11): p. 3887–3895.

27. Pardoll, D.M., The blockade of immune checkpoints in cancer immunotherapy. Nature reviews cancer, 2012. 12(4): p. 252–264.

28. Sharma, P., et al., Primary, adaptive, and acquired resistance to cancer immunotherapy. Cell, 2017. 168(4): p. 707–723.

29. Liu, J., et al., Glycosylation and Its Role in Immune Checkpoint Proteins: From Molecular Mechanisms to Clinical Implications. Biomedicines, 2024. 12(7): p. 1446.

30. Agostini, M., P. Traldi, and M. Hamdan, Proteomic Investigation of Immune Checkpoints and Some of Their Inhibitors. International Journal of Molecular Sciences, 2024. 25(17): p. 9276.

31. Kuol, N., et al., PD-1/PD-L1 in disease. Immunotherapy, 2018. 10(2): p. 149–160.

32. Shiravand, Y., et al., Immune checkpoint inhibitors in cancer therapy. Current Oncology, 2022. 29(5): p. 3044–3060.

33. Liang, S.C., et al., Regulation of PD-1, PD-L1, and PD-L2 expression during normal and autoimmune responses. European journal of immunology, 2003. 33(10): p. 2706–2716.

34. Lin, X., et al., Regulatory mechanisms of PD-1/PD-L1 in cancers. Molecular Cancer, 2024. 23(1): p. 108.

35. Han, Y., D. Liu, and L. Li, PD-1/PD-L1 pathway: current researches in cancer. American journal of cancer research, 2020. 10(3): p. 727.

36. Iwai, Y., et al., Involvement of PD-L1 on tumor cells in the escape from host immune system and tumor immunotherapy by PD-L1 blockade. Proceedings of the National Academy of Sciences, 2002. 99(19): p. 12293–12297.

37. Chen, G., et al., Transdermal cold atmospheric plasma-mediated immune checkpoint blockade therapy. Proceedings of the National Academy of Sciences, 2020. 117(7): p. 3687–3692.

38. Van Der Spoel, D., et al., GROMACS: fast, flexible, and free. Journal of computational chemistry, 2005. 26(16): p. 1701–1718.

39. Abraham, M.J., et al., GROMACS: High performance molecular simulations through multi-level parallelism from laptops to supercomputers. SoftwareX, 2015. 1: p. 19–25.

40. Berendsen, H.J., D. van der Spoel, and R. van Drunen, GROMACS: A message-passing parallel molecular dynamics implementation. Computer physics communications, 1995. 91(1-3): p. 43–56.

41. Berman, H.M., et al., The protein data bank. Acta Crystallographica Section D: Biological Crystallography, 2002. 58(6): p. 899–907.

42. Margreitter, C., D. Petrov, and B. Zagrovic, Vienna-PTM web server: a toolkit for MD simulations of protein post-translational modifications. Nucleic acids research, 2013. 41(W1): p. W422–W426.

43. Takai, E., et al., Chemical modification of amino acids by atmospheric-pressure cold plasma in aqueous solution. Journal of Physics D: Applied Physics, 2014. 47(28): p. 285403.

44. Roither, B., C. Oostenbrink, and W. Schreiner, Molecular dynamics of the immune checkpoint programmed cell death protein I, PD-1: conformational changes of the BC-loop upon binding of the ligand PD-L1 and the monoclonal antibody nivolumab. BMC bioinformatics, 2020. 21: p. 1–12.

45. Schmid, N., et al., Definition and testing of the GROMOS force-field versions 54A7 and 54B7. European biophysics journal, 2011. 40: p. 843–856.

46. Berg, J., Calculating near-densest lattice packings of non-convex objects to minimize computational box volumes in Molecular Dynamics simulations. 2003, Faculty of Science and Engineering.

47. Pfeiffer, S., D. Fushman, and D. Cowburn, Impact of Cl− and Na+ ions on simulated structure and dynamics of βARK1 PH domain. Proteins: Structure, Function, and Bioinformatics, 1999. 35(2): p. 206–217.

48. Mark, P. and L. Nilsson, Structure and dynamics of the TIP3P, SPC, and SPC/E water models at 298 K. The Journal of Physical Chemistry A, 2001. 105(43): p. 9954–9960.

49. Humphrey, W., A. Dalke, and K. Schulten, VMD: visual molecular dynamics. Journal of molecular graphics, 1996. 14(1): p. 33–38.

50. DeLano, W.L., Pymol: An open-source molecular graphics tool. CCP4 Newsl. Protein Crystallogr, 2002. 40(1): p. 82–92.

